# NK cells acquire epigenetic memory of LPS-induced systemic inflammation

**DOI:** 10.1101/506014

**Authors:** Orhan Rasid, Christine Chevalier, Tiphaine Camarasa, Catherine Fitting, Jean-Marc Cavaillon, Melanie Anne Hamon

## Abstract

Natural killer cells are unique mediators of innate immunity, and as such, an attractive target for immunotherapy. Following viral infection, NK cells display immune memory properties, defined by heightened responses to re-stimulation, an expansion of specific NK cell sup-populations and a protective role against re-infection. However, similar memory to bacterial infection or systemic inflammation, and the molecular mechanisms behind NK cell memory remain elusive. Here we show that following LPS-induced endotoxemia in mice, NK cells acquire cell-intrinsic memory properties as displayed by an amplified production of IFNγ upon secondary stimulation. NK cell memory is acquired even under the post-endotoxemic suppressive environment and is detectable for at least 9 weeks. Furthermore, we define an epigenetic mechanism essential for NK cell memory, where an H3K4me1-marked latent enhancer is uncovered at the *ifng* locus. Chemical inhibition of histone methyltransferase activity erased the enhancer and prevented NK cells from acquiring memory. Thus, NK cells develop memory to LPS during endotoxemia, in a histone methylation-dependent manner, which ensures a heightened response to secondary stimulation and confers protection against bacterial infection.

## INTRODUCTION

In recent years, accumulating evidence of memory responses mediated by innate immune cells have blurred the boundaries between innate and adaptive immunity(*1, 2*). Monocytes and macrophages have been described to mediate a type of innate immune memory, termed trained immunity(*3*). Briefly, this refers to the long-lasting altered responsiveness of cells to a secondary stimulation, with either lower (tolerized) or higher (trained) inflammatory responses, involving epigenetic modifications and metabolic rewiring(*4*). While trained immunity in phagocytes is non-specific and results in a general hyper or hypo-responsiveness to re-stimulation, other innate cells such as NK cells can exhibit a high degree of antigen specificity in a secondary response(*3, 5*). NK cells have been shown to display memory properties to viral infection, cytokine stimulation or hapten-induced contact-hypersensitivity, but evidence for bacterial infection remains scarce(*5*). The specificity of the recall response together with proliferation of specific receptor-defined sub-populations make NK cell memory more akin to classical adaptive memory mediated by T cells(*6*).

NK cell memory was demonstrated in several *in vivo* models of viral infection, particularly mouse cytomegalovirus (MCMV), where the murine Ly49H receptor, which specifically recognize the MCMV m157 glycoprotein, defines the population of responsive NK cells that acquire memory(*7*). In humans infected with CMV the NKG2C receptor was involved in ensuring NK cell responses to specific viral peptides(*8*). At the molecular level, NK cell memory to CMV has been correlated with modified chromatin states(*9*). Mouse and human memory-like NK cells have modulated DNA methylation levels, which are notably reduced at the *ifng/IFNG* gene locus(*10*), and more accessible chromatin at effector genes as assessed by ATAC-Seq(*11*). However, whether these or other epigenetic marks are important for establishing and maintaining memory, remains to be determined.

A common feature of all NK cell memory models is their dependence on an initial inflammatory response with several key cytokines implicated as critical factors for NK cell memory acquisition(*6*). Systemic inflammation, such as sepsis or severe trauma, represents one of the most complex paradigms in immunology and involves an overzealous inflammatory response that often represents a life-threatening condition(*12*). NK cells are key players and one of the main drivers of systemic inflammation during the acute phase, but following, they are severely incapacitated, similarly to monocytes and T cells(*13–15*). Specifically, NK cells have impaired cytotoxicity and cytokine production in mouse models and human sepsis patients(*16–19*). Altogether, these immune consequences are known as the “immunosuppressive” phase of systemic inflammation(*20*), a state of immune dysfunction that is thought to persist beyond the acute event, causing poor quality of life and increased susceptibility to infection(*21*). However, while short-term consequences on the immune system are fairly well explored, the long-term effects remain understudied. Indeed, the long-term status of NK cells after systemic inflammation remains to be explored and could have important implications, given their well-known role as critical effectors in viral and bacterial infections(*22*).

Herein, we used an endotoxemia model to explore the lasting impact of systemic inflammation on NK cells. We found that NK cells acquire cell-intrinsic memory properties, demonstrated by increased IFNγ production upon re-challenge, up to 9 weeks after endotoxemia. Strikingly, systemic inflammation revealed a latent enhancer at the *ifng* locus in memory NK cells and blocking this, along with histone methylation, prevented acquisition of memory. Thus, we report that despite the suppressive immune environment that follows systemic inflammation, NK cells retain memory properties that are epigenetically encoded and help protect mice against bacterial infection.

## RESULTS

### 1. NK cells are responsive after endotoxemia and acquire memory properties despite being in a suppressive environment

In order to assess the long-term effects of systemic inflammation on NK cells, we used an endotoxemia model (10 mg/kg LPS, i.p.) causing acute inflammation but low mortality, with full clinical recovery by day 7 (Fig. 1a,b,c). Indeed, mice exhibited transient weight loss, clinical signs of inflammation and limited morbidity. Using this model, we had previously shown that NK cells are systemically activated over a 48-hour acute period(*23*). In the current study, we focused on the impact of endotoxemia on NK cells, 14 days after LPS injection. First, we phenotypically defined NK cells at 14 days after endotoxemia (full gating strategy in fig. S1). We found no significant differences in NK cell percentages or total numbers in the spleen of mice that received PBS injections as controls (D14PBS) or experienced endotoxemia 14 days prior (D14LPS) (Fig. 1d,e). Furthermore, although we found some variations in NK cell phenotype (fig. S1b,c,d), such differences did not correlate with an altered baseline of IFNγ expression detected upon *ex-vivo* staining in D14LPS mice compared to D14PBS (fig. S1e). Therefore, altogether these data indicate that 14 days after endotoxemia, while NK cells acquire slight phenotypical differences, they return to a resting state, similar to that of naïve cells.

**Fig. 1.**
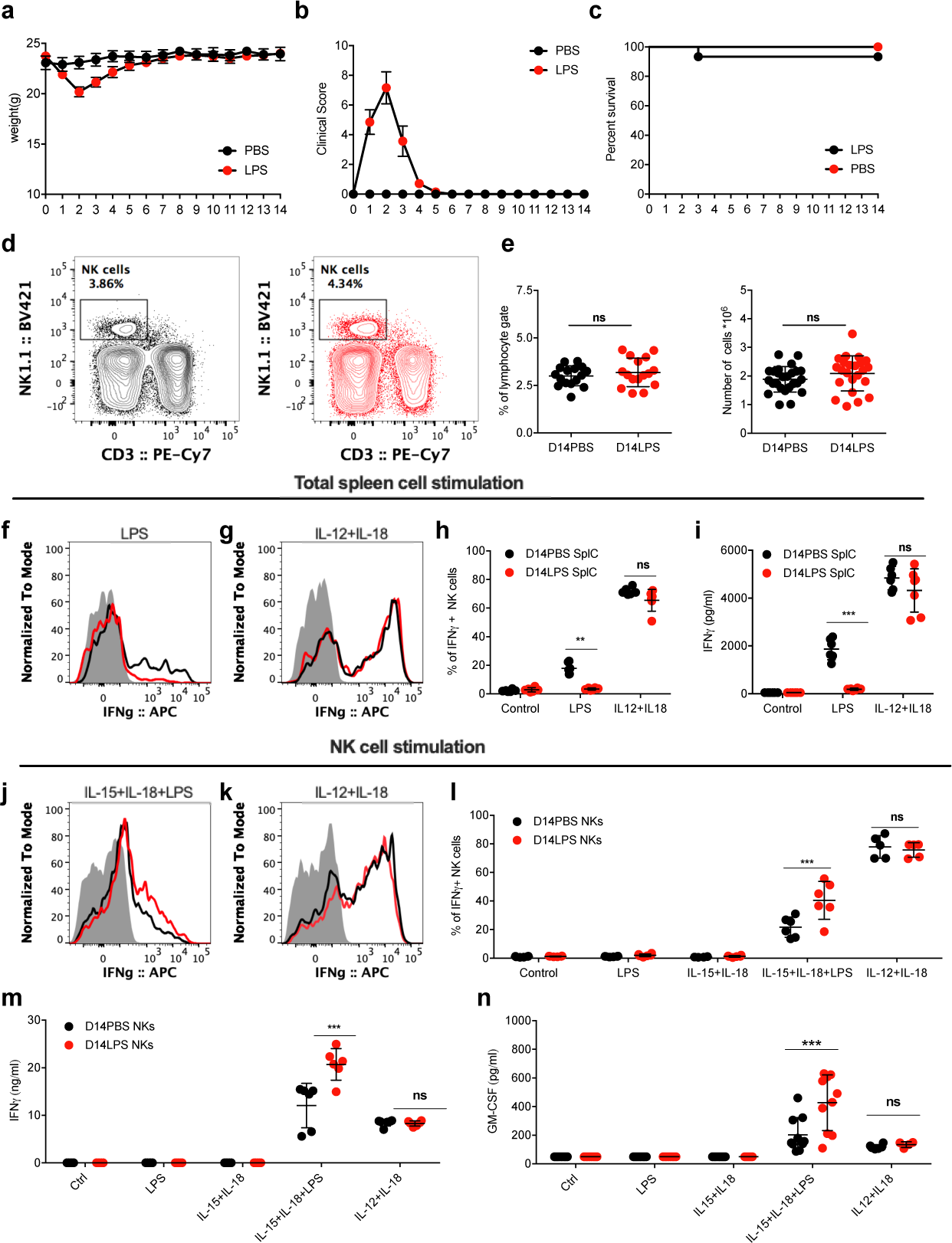
NK cells acquire memory-like features 14 days after endotoxemia. C57BL/6 mice were injected with either PBS or LPS (10 mg/kg) intraperitoneally and monitored 14 after which they were used to assess NK cell status *ex vivo*. **a)** Weight, **b)** Clinical score and **c)** Survival monitoring following injection of PBS or LPS. Spleen NK cells from D14PBS and D14LPS mice were analyzed: **d)** Representative gating for NK cells in both conditions **e)** Percentages of NK cells among lymphocytes and total NK cell numbers. Splenocytes (SplC) from D14PBS and LPS mice were stimulated *in vitro* with cytokines (10ng/ml) or LPS (100ng/ml) overnight. Cell were stained for flow cytometry analysis to assess levels of intracellular IFNγ and supernatants were collected for cytokine ELISAs. **f-g)** Representative overlay histograms and **h)** quantification of IFNγ expression by NK cells in respective conditions. **i)** Total IFNγ levels in supernatants from SplC. NK cells were enriched from spleens of D14PBS and LPS mice and stimulated *in vitro* with cytokines and/or LPS overnight. **j-k)** Representative overlay histograms and **l)** quantification of IFNγ expression by NK cells in respective conditions. Total **m)** IFNγ and **n)** GM-CSF levels in supernatants from NK cells in respective conditions. Dots represent individual mice. Data represent one out of least 3 repeats with n≥ 5 mice/group. ns, not significant, *p<0.05, **p<0.01, ***p<0.001. Mann–Whitney test comparing D14PBS and D14LPS cell values.

We next tested the responsiveness of NK cells from post-endotoxemia mice *in vitro* both within the environment of the spleen and separately as purified cells. We first stimulated total spleen cells with LPS and observed significantly reduced IFNγ positive NK cell percentages from D14LPS samples as compared to D14PBS controls, as well as reduced total IFNγ secretion in supernatants (Fig. 1f,h,i). In correlation with the lower levels of NK cell activation in D14LPS mice, we found a significant increase in percentages of regulatory T cells (Tregs) and accumulation of immature myeloid cells in these mice as compared to D14PBS controls (fig. S2). Therefore, the immune environment in the spleen 14 days after systemic inflammation is suppressive and affects NK cells, which is in agreement with previous reports(*24–26*). Interestingly, stimulation with IL-12+IL-18 stimulation, targeted NK cell activators, induced similar levels of NK cells response between D14PBS and D1LPS (Fig. 1g,h,i). These results were surprising and suggested that while post-endotoxemia spleen cells cannot activate NK cells in response to LPS, the intrinsic responsiveness of NK cells, at least to IL-12+IL-18, was not altered 14 days after endotoxemia.

To further study the response of NK cells independently of the splenic environment, we stimulated enriched NK cells (~80%) *in vitro*. We had previously shown that isolated NK cells respond to LPS stimulation *in vitro* if co-stimulated with the cytokines IL-15+IL-18, which do not induce sustained activation on their own(*16*). In agreement, while overnight stimulation with IL-15+IL-18 induced no detectable response, IL-12+IL-18 stimulation induced a sustained IFNγ production. However, no differences were observed between control D14PBS and D14LPS NK cells in terms of IFNγ+ percentages or total IFNγ production in supernatants (Fig. 1k,l,m), as we had observed when stimulating total spleen cells. In contrast, upon stimulation with IL-15+IL-18+LPS, we found that D14LPS NK cells had significantly higher percentages of IFNγ+ cells compared to control D14PBS NK cells (Fig. 1j,l). The higher responsiveness of D14LPS NK cells was confirmed by measuring total levels of secreted IFNγ, as well as GM-CSF, in cell culture supernatants (Fig. 1m,n). Altogether, these results indicate that following systemic inflammation, NK cells have a significantly increased responsiveness to LPS *in vitro*, a memory-like property.

Since our *in vitro* stimulations are performed on enriched NK cells, we cannot exclude a possible effect of contaminants such as myeloid or T cells. To exclude this possibility, we performed the same experiments as above, but mixed enriched (~80%) naïve CD45.1 NK cells with CD45.2 D14PBS or D14LPS NK cells, which were highly purified (~98%). In this way, each well containing D14PBS or D14LPS NK cells has an internal control of naïve NK cells, which are subjected to the same influence of possible contaminants. Cocultured naïve and D14PBS NK cells responded similarly to IL-15+IL-18+LPS stimulation, as expected (fig. S1f,g). Furthermore, naïve cells cocultured with D14LPS NK cells reached slightly higher levels of IFNγ expression compared to D14PBS NK cells but these were significantly surpassed by the cocultured D14LPS NK cells (fig. S1g,h). Therefore, these results argue against a major effect of accessory cell contaminants in our *in vitro* stimulation system and strongly support the cell-intrinsic nature of memory-like responses observed in D14LPS NK cells. Notably, the unaltered production of IFNγ in response to cytokine stimulation would suggest that changes in NK cell reactivity after endotoxemia do not simply reflect a state of hyperresponsiveness. These observations are in agreement to what was previously shown for memory NK cells(*27*). Taken together, our results show that 2 weeks following endotoxemia, NK cells are still responsive and have acquired cell-intrinsic memory properties, while being in a suppressive environment that restrains their activation during a secondary challenge with LPS.

### 2. Endotoxemia-induced NK cell memory is detectable *in vivo* and is cell-intrinsic

We further explored the memory responses of NK cells *in vivo* upon LPS challenge. In order to compare D14LPS NK cells to naïve counterparts in the same immune environment, we used an adoptive transfer model in which naïve congenic NK cells were transferred into post-endotoxemic mice (Fig. 2a). We transferred naïve CD45.1 NK cells into CD45.2 mice that had been treated with PBS or LPS 13 days prior. On day 14, recipient mice were re-challenged with PBS or LPS and NK cell responses in the spleen were assessed 6 hours later by flow-cytometry. Transferred naïve NK cells were readily identified by CD45.1 staining and their IFNγ expression was quantifiable (Fig. 2b). Importantly, control PBS injection induced no detectable IFNγ in neither the endogenous or transferred NK cell populations, further demonstrating that 14 days after endotoxemia, NK have returned to a resting state (Fig. 2c). LPS re-challenge induced significant expression of IFNγ by endogenous NK cells as well as transferred naïve NK cells in D14PBS mice (Fig. 2d, left). Since both endogenous and transferred NK cells responded at similar levels, the transfer procedure itself had no influence on IFNγ response. In D14LPS mice, the percentages of IFNγ+ NK cells were significantly lower as compared to those from D14PBS mice (Fig. 2d), as expected given the suppressive environment we described. Strikingly, endogenous D14LPS NK cells had significantly higher percentages of IFNγ+ than transferred naïve NK cells (Fig. 2d, right). These results demonstrate that NK cell memory acquired after endotoxemia results in increased responsiveness to re-stimulation *in vivo* even under the control of the suppressive environment induced by systemic inflammation.

**Fig. 2.**
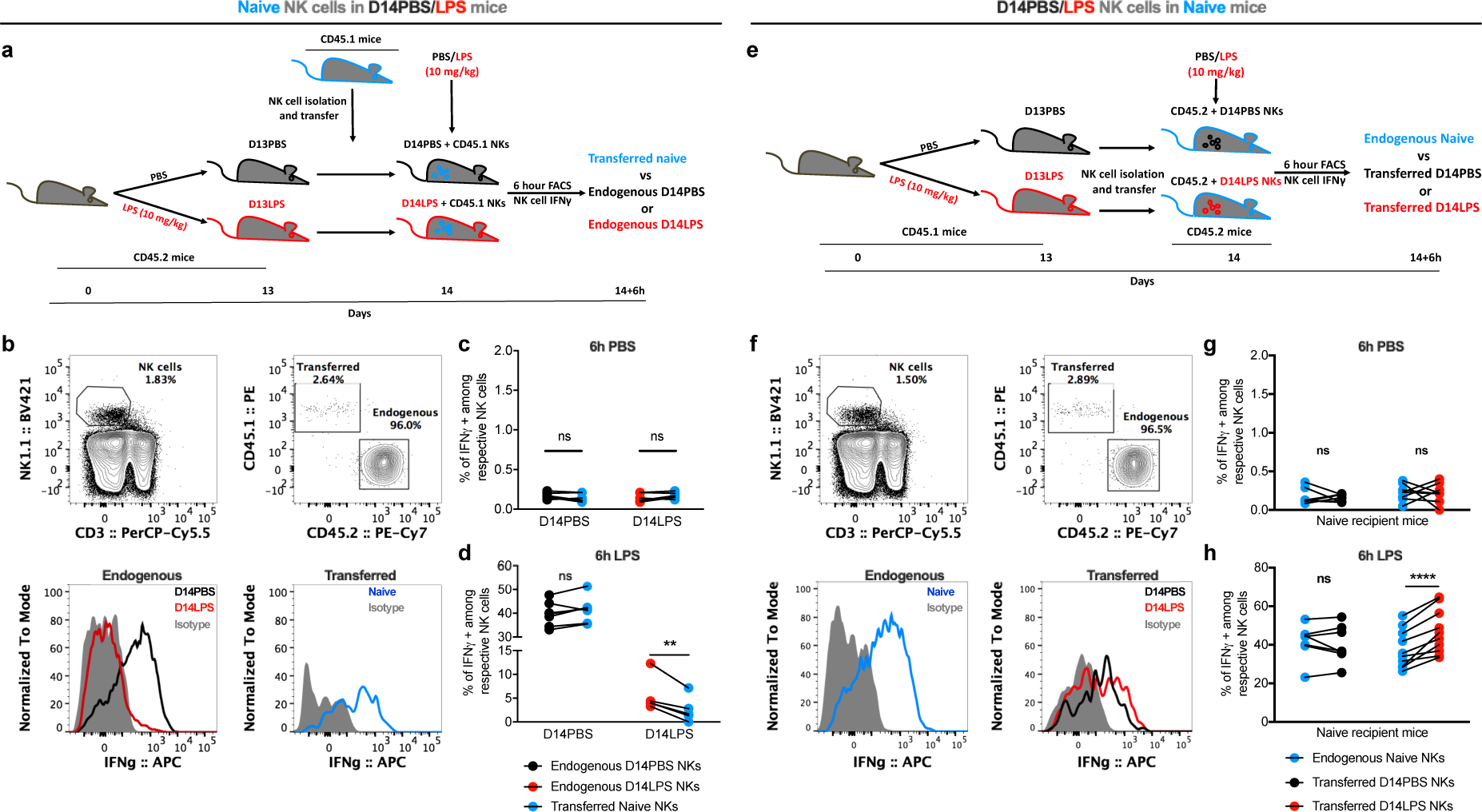
NK cells retain cell-intrinsic memory properties *in vivo* after endotoxemia under a suppressive environment. NK cells enriched from spleens of naïve CD45.1 mice were transferred into CD45.2 mice (1×10^6^ cells/recipient, intravenously) injected 13 days before with PBS (D13PBS) or LPS (10mg/kg, i.p.) (D13LPS). Recipient mice were challenged one day later with PBS or LPS (10mg/kg) and sacrificed 6 hours later. Splenocytes were processed for flow cytometry to levels of intracellular IFNγ in endogenous vs transferred NK cells. **a)** Experimental transfer scheme **b)** Representative gating strategy **c)** Data summary grouped before-after plots depicting percentages of IFNγ+ cells among endogenous CD45.2 D14PBS NK cells (black dots), transferred CD45.1 naive NK cells (blue dots) and endogenous CD45.2 D14LPS NK cells (red dots) at 6 hours after PBS or **d)** LPS rechallenge. NK cells were enriched from spleens of CD45.1 mice injected 13 days before with PBS (D13PBS) or LPS (10mg/kg, i.p.) (D13LPS) and were transferred into naïve CD45.2 mice (1×10^6^ cells/recipient, intravenously). Recipient mice were challenged and samples harvested as above. **e)** Experimental transfer scheme **f)** Representative gating strategy **g)** Data summary grouped before-after plots depicting percentages of IFNγ+ cells among endogenous CD45.2 naive NK cells (blue dots), transferred D14PBS CD45.1 NK cells (black dots) and transferred D14LPS CD45.2 NK cells (red dots) at 6 hours after PBS or **h)** LPS re-challenge. Dots represent individual mice. Data are representative of one experiment out of 3 repeats with n≥ 5 mice/group. ns, not significant, * p<0.05. Wilcoxon paired test comparing D14Sln, D14LPS and naïve cell values under respective conditions.

The Ly49H and NKG2C receptors have both been implicated in mediating NK cell memory responses to MCMV and CMV, respectively(*7, 28*), and we observed significant differences in the proportions of NK cell subpopulations defined by these receptors after endotoxemia (fig. S1c). We thus investigated whether subpopulations defined by these receptors might be involved in mediating memory in our model. We compared populations defined by NKG2A/C/E and Ly49H in both D14PBS and D14LPS mice, and found no difference in the percentages of IFNγ+ cells upon LPS re-challenge between positive or negative populations (fig. S3). These results suggest that neither of these subpopulations is involved in the observed results, thus, we conducted the rest of our study using the bulk NK cell population. To explore memory NK cell function independently of the surrounding immunological environment, we turned to another adoptive transfer system in which we can evaluate intrinsic NK cell properties. For this, we transferred D13PBS or D13LPS NK cells into naïve congenic mice one day before re-challenge with PBS or LPS (Fig. 2e). Thus, CD45.2 D14PBS or D14LPS NK cells were compared to naïve endogenous CD45.1 NK cells 6 hours after PBS or LPS challenge (Fig. 2f). PBS injection revealed no differences in basal IFNγ expression between transferred CD45.2 and endogenous naive CD45.1 NK cells, suggesting that both D14PBS or D14LPS NK cells remain in a resting state even outside the suppressive environment of the post-endotoxemia spleen (Fig. 2g). Upon LPS injection, a high percentage of IFNγ+ cells is detected both in control transferred D14PBS NK cells and endogenous naïve NK cells, at similar values (Fig. 2h, left). Strikingly, significantly higher percentages of IFN+ cells were detected among the transferred D14LPS NK cells compared to the respective endogenous NK cells (Fig. 2h, right). These results firmly demonstrate that NK cells intrinsically acquire memory features 14 days following systemic inflammation and this is independent of the surrounding immune environment.

### 3. Endotoxemia-induced NK cell memory persists for at least 9 weeks

The lasting potential is an important feature of a memory response, which we explored 9 weeks after endotoxemia. For this we tested the responsiveness of NK cells by performing adoptive transfer schemes similar to those in Fig. 2a and e, but 9 weeks after PBS or LPS injection (We9PBS/LPS). Remarkably, memory properties of NK cells were retained at this moment. Indeed, endogenous We9LPS NK cells responded with more IFNγ than transferred naïve NK cells, despite the suppressive environment, which was still present at this time (Fig. 3a), albeit less restrictive than at day 14. Additionally, transferred We9LPS NK cells still surpassed endogenous NK cells in IFNγ+ percentages in naïve recipient mice (Fig. 3b). We also performed *in vitro* stimulations of enriched NK cells and observed We9LPS NK cells still being significantly more responsive to IL-15+IL-18+LPS that We9PBS NK cells (Fig. 3c,d). Therefore, post-endotoxemia NK cells retain memory properties, characterized by an increased IFNγ production to LPS challenge, a phenotype that persists for at least 9 weeks.

**Fig. 3.**
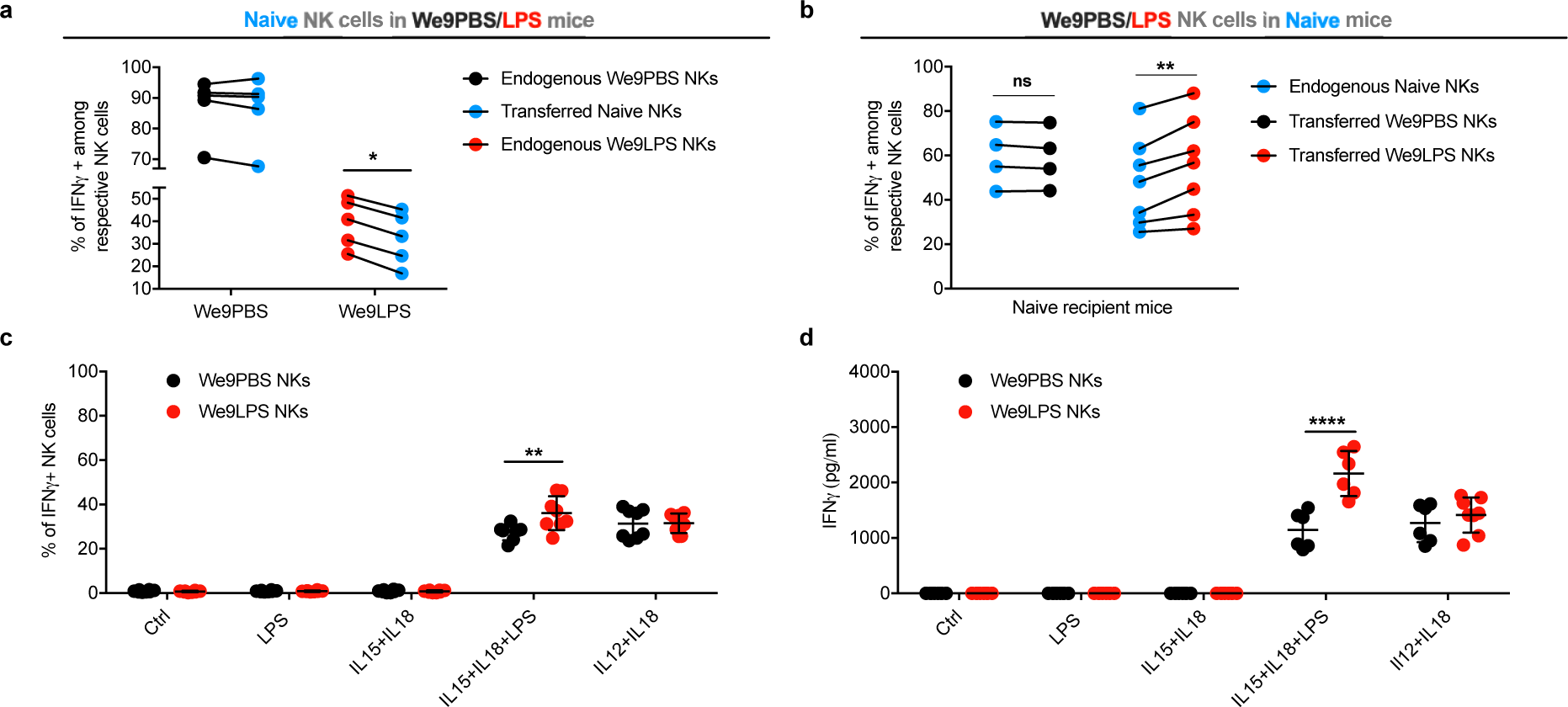
NK cells retain memory properties for at least 9 weeks after endotoxemia. NK cells enriched from spleens of naïve mice were transferred into congenic mice (1×10^6^ cells/recipient, intravenously) injected 9 weeks before with PBS (We9PBS) or LPS (10mg/kg, i.p.) (We9LPS). Recipient mice were challenged one day later with LPS (10mg/kg) and sacrificed 6 hours later. Splenocytes were processed for flow cytometry to levels of intracellular IFNγ in endogenous vs transferred NK cells. **a)** Data summary grouped before-after plots depicting percentages of IFNγ+ cells among endogenous We9PBS NK cells (black dots), transferred naive NK cells (blue dots) and endogenous We9LPS NK cells (red dots) at 6 hours after LPS rechallenge. NK cells were enriched from spleens of mice injected 9 weeks before with PBS (We9PBS) or LPS (10mg/kg, i.p.) (We9LPS) and were transferred into naïve congenic mice (1×10^6^ cells/recipient, intravenously). Recipient mice were challenged and samples harvested as above. **b)** Data summary grouped before-after plots depicting percentages of IFNγ+ cells among endogenous naive NK cells (blue dots), transferred We9PBS NK cells (black dots) and transferred We9LPS NK cells (red dots) at 6 hours after LPS rechallenge. NK cells were enriched from spleens of We9PBS and LPS mice and stimulated *in vitro* with cytokines (10ng/ml) or LPS (100ng/ml) overnight. Quantification of **c)** IFNγ expression by NK cells and total levels of **d)** IFNγ in respective conditions. Dots represent individual mice. Data are representative of one experiment out of 3 repeats with n≥4 mice/group. ns, not significant, *** p<0.001. Wilcoxon paired test comparing D14Sln, D14LPS and naïve cell values under respective conditions (a-b) and Mann– Whitney test comparing D14PBS and D14LPS values (c-d).

### 4. Memory cells derive from NK cells that experienced endotoxemia

To investigate the origin of D14LPS NK cells and whether these cells are the same (or descendants of the same) cells present at the time of endotoxemia, or new cells that matured and arrived in the spleen after systemic inflammation, we made use of the adoptive transfer system described in Fig. 4a. In this system, traceable congenic NK cells are transferred into recipient mice 1 day prior to endotoxemia and are followed through day 14. An additional transfer of naïve NK cells is performed 7 days after endotoxemia in order to have control naïve cells at day 14 in the same recipient mouse. Thus, we have a model in which a defined population of pre-transferred CD45.1 NK cells will have been exposed to the same control (D14PBS) or inflammatory (D14LPS) conditions as the endogenous CD45.1/2 NK cells, and a post-transferred population of naïve CD45.2 NK cells that will serve as additional naïve controls. Using this model, we re-challenged D14PBS and D14LPS CD45.1/2 mice with PBS or LPS and assessed NK cell IFNγ expression. Due to the time period between the pre-transfer and the response readout, and the potential for NK cells plasticity of NK cells into type 1 innate lymphoid cells (ILC1s) in inflammatory and immunosuppressive environments like tumors(*29*), we increased the stringency of our gating strategy. We introduced an additional marker to our panel, and defined our NK cells as CD3^-^ NK1.1^+^CD49b^+^ (Fig. 4b), thereby excluding interference of ILC1s in these experiments. Pre-transferred CD45.1 and post-transferred CD45.2 NK cells were readily detectable. Upon PBS injection, we observed no difference in basal IFNγ expression by any NK cell population in D14PB or LPS mice (Fig. 4c). Furthermore, upon LPS injection into D14PBS mice, we observed similar percentages of IFNγ+ cells among the pre-transferred D14PBS CD45.1, endogenous CD45.1/2 D14PBS, and post-transferred naïve CD45.2 NK cell populations (Fig. 4d, left and fig. S4a), demonstrating that in the absence of prior inflammation, transferred and endogenous cell populations behave similarly. However, upon LPS injection into D14LPS mice, we observe an increase in percentages of IFNγ+ cells in endogenous D14LPS CD45.1/2 NK cells compared to post-transferred naïve CD45.2 NK cells (Fig. 4d, right and fig. S4b). Most interestingly, the pre-transferred D14LPS CD45.1 NK cells also showed significantly higher IFNγ+ percentages compared to post-transferred naïve CD45.2 NK cells and similar levels to endogenous cells (Fig. 4d, right). This transfer system was also the ideal approach to address the involvement of NK cell-intrinsic TLR signaling in our endotoxemia model. Indeed, *in vitro* testing is hampered by the fact that a chronic lack of tonic TLR signaling lowers NK cell responsiveness in general, as indicated in literature(*30*). By pre-transferring TLR2/4KO NK cells, we found that both primary and memory responses of NK cells during LPS challenge were independent of TLR2 and 4 (fig S4c). Altogether, these results demonstrate that NK cells present at the moment of systemic inflammation, or descendants of the original cells, acquire memory properties after endotoxemia, in a TLR2 and −4-independent manner.

**Fig. 4.**
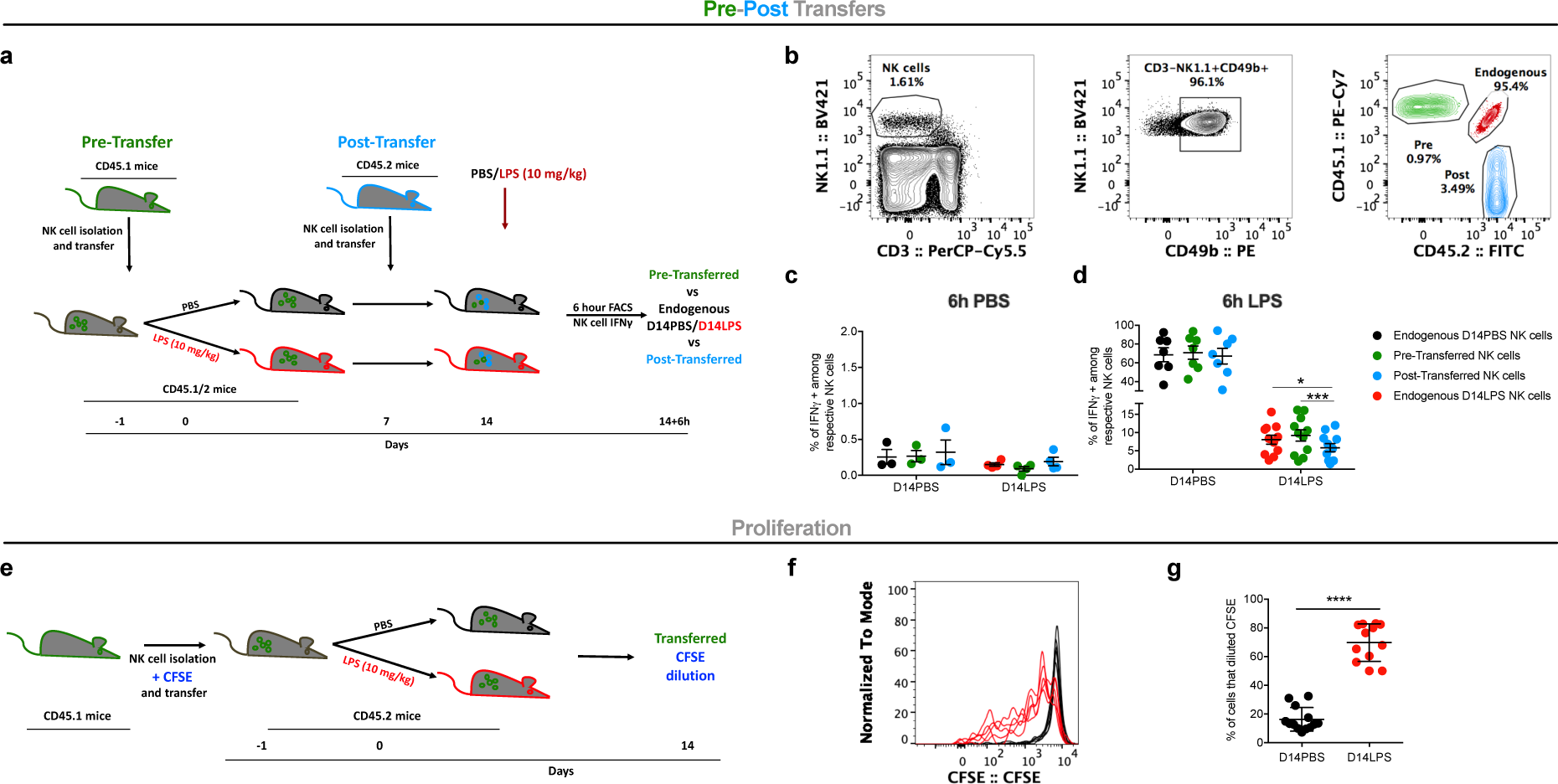
Memory-like cells derive from NK cells that experienced endotoxemia. NK cells enriched from spleens of CD45.1 mice were Pre-Transferred into CD45.1/2 mice (1×10^6^ cells/recipient, intravenously), one day before recipients were injected with PBS or LPS (10mg/kg, i.p.). Naïve CD45.2 enriched NK cells were Post-Transferred into the CD45.1/2 mice, 7 days after PBS/LPS injection. On day 14 with CD45.1/2 D14PBS and D14LPS mice were rechallenged with LPS and sacrificed 6 hours later to assess intracellular IFNγ levels in endogenous, pre- and post-transferred NK cells. **a)** Experimental transfer scheme **b)** Representative gating strategy **c)** Data summary scatter plots depicting percentages of IFNγ+ cells among endogenous CD45.1/2 D14PBS NK cells (black dots), Pre-Transferred CD45.1 NK cells (green dots), Post-Transferred CD45.2 (blue dots) and endogenous CD45.1/2 D14LPS NK cells (red dots) at 6 hours after PBS or **d)** LPS re-challenge. Enriched CD45.1 naïve NK cells were labeled with CFSE and transferred into CD45.2 mice one day before PBS or LPS (10mg/kg, i.p.) injection. On day 14, spleens were harvested from D14PBS and D14LPS mice, and stained for transferred NK cell identification in order to assess CFSE dilution. **e)** Experimental transfer scheme **f)** Representative overlay histogram showing CFSE dilution of CD45.1 transferred cells in D14PBS (black lines) and D14LPS (red lines). **g)** Data summary showing percentages of cells that diluted CFSE (gated on all cells before the highest intensity peak). Dots represent individual mice. Data are representative of one experiment out of 3 repeats with n≥ 5 mice/group. ns, not significant, * p<0.05, *** p<0.001. Wilcoxon paired test comparing D14Sln, D14LPS and naïve cell values under respective conditions.

To determine whether the endotoxemia-induced memory cells have proliferated during the course of 14 days, we pre-transferred CFSE-labeled CD45.1 NK cells into CD45.2 mice followed by PBS/LPS injection (Fig. 4e). At day 14, in D14PBS mice we observed minimal CFSE dilution of pre-transferred NK cells (Fig. 4f). In contrast, in D14LPS mice, the majority of pre-transferred NK had divided as indicated by several peaks of decreasing CFSE signal (Fig. 4f,g). Our results thus demonstrate that D14LPS memory cells derive from NK cells that were exposed to systemic inflammation and maintain memory properties after undergoing several rounds of proliferation.

### 5. Histone methylation at an *ifng* enhacer is involved in NK cell memory

Since innate immune memory in macrophages and NK cells has been proposed to rely on epigenetic modifications, we explored whether this might be the case for post-endotoxemia memory NK cells. To investigate the possible epigenetic mechanism responsible from memory features we studied the histone marks associated with upstream regulatory regions of the *ifng* locus by chromatin immunoprecipitation followed by qPCR (ChIP-qPCR). We probed the *ifng* promoter and several possible enhancer regions, which had been previously described as being crucial for IFNγ transcription(*31*). As positive and negative controls, we probed ActB, a housekeeping gene known to be expressed and Hoxc8, that is silent in hematopoietic cells, respectively (Fig. 5a). We focused on H3K4me1, a histone modification that marks active enhancers. Interestingly, we identified one region of the *ifng* locus, located −22 kb from the transcriptional start site, that showed significant enrichment for H3K4me1 in D14LPS NK cells (Fig. 5b). The same region was not marked with H3K4me1 in D14PBS NK cells. Therefore, endotoxemia reveals a latent enhancer at the *ifng* locus, thereby poising NK cells for increased IFNγ expression. Such results strongly suggest that post-endotoxemia NK cell memory has an epigenetic component.

**Fig. 5.**
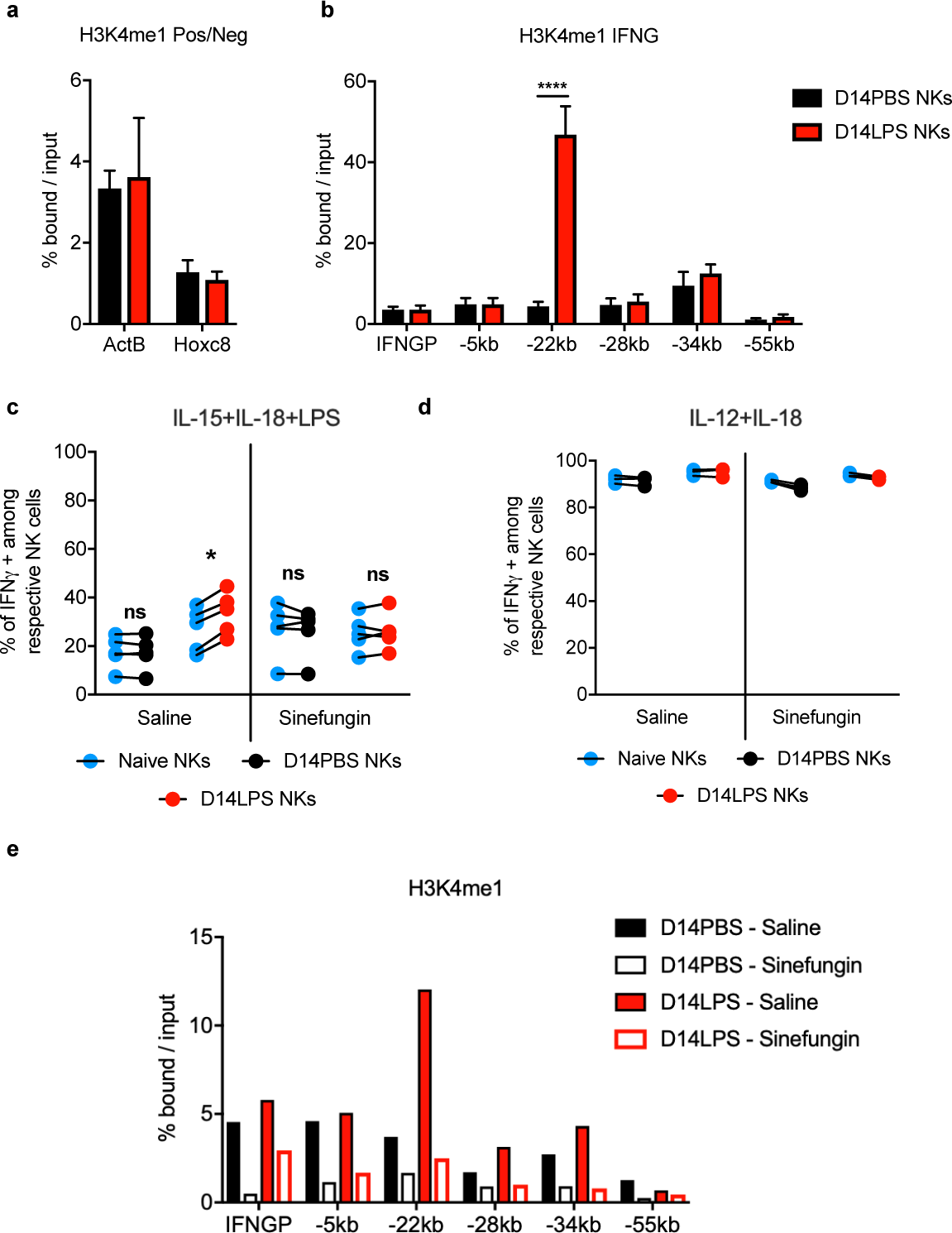
Histone methylation maintains NK cell memory after endotoxemia. NK cells were isolated from spleens of mice injected 14 days before with PBS (D14PBS) or LPS (10mg/kg, i.p.) (D14LPS). Approximately 3×10^6^ highly purified NK cells were fixed and chromatin was extracted and sheared. ChIP for H3K4me1 and H3K4me3 was perfomed and resulting positive and input fractions of the chromatin were amplified by PCR for the indicated targets. **a)** Enrichment percentage for H3K4me1 pulldown on positive (ActB) and negative (Hox8) controls. **b)** Enrichment percentage for H3K4me1 pulldown on different regions of the *ifng* locus. **c** and **d)** Mice were injected with PBS or LPS (10mg/kg, i.p.). On day 2, mice received either Saline or Sinefungin (10mg/kg). On day 14, NK cells were higly purified from mice D14PBS or D14LPS NK cells that were treated with Saline or Sinefungin. NK cells from each group were mixed with enriched naïve congenic NK cells and stimulated overnight in the indicated conditions. Cells were stained for flow cytometry analysis of intracellular IFNγ levels. Data summary grouped before-after plots depicting percentages of IFNγ+ cells among naïve (blue dots), D14PBS (black dots) and D14LPS (red dots) NK cells. Treatment groups underneath graph. **e)** ChIP for H3K4me1 on chromatin of NK cells purified from D14PBS-Saline, D14PBS-Sinefungin, D14LPS-Saline, D14LPS-Sinefungin mice. Enrichment percentage for H3K4me1 pulldown on different regions of the *ifng* locus for each group is showed. Data are representative of one experiment out of at least 2 repeats with n≥3 mice/group except for panel d which was performed once. ns, not significant, *p<0.05,*** p<0.001. Mann–Whitney test (a,b) and Wilcoxon paired test (c) comparing D14Sln, D14LPS and naïve cell values under respective conditions.

To test the involvement of histone methylation on memory acquisition by NK cells after endotoxemia, we used a chemical methyltransferase inhibitor, sinefungin, previously reported to reduce H3K4me1 *in vivo*(*32*). We treated mice with sinefungin or saline, as a vehicle control, 2 days after PBS or LPS injection. At day 14 we purified NK cells from the 4 groups of mice (D14PBS - Saline, D14PBS - Sinefungin, D14LPS - Saline and D14LPS - Sinefungin) and stimulated them with IL-15+IL-18+LPS in co-culture with congenic naïve NK cells. Strikingly, while saline treated D14LPS NK cells showed memory level responses, with higher IFNγ+ percentages, as compared to the co-cultured naïve NK cells and D14PBS NK cells, sinefungin treated D14LPS NK cells lost memory properties, responding at similar levels to co-cultured naïve NK cells (Fig. 5c). Importantly, sinefungin treatment did not block the response of D14PBS or D14LPS NK cells to IL-12+IL-18 (Fig. 5d). These results, demonstrate that methyltransferase-dependent modifications in D14LPS NKs are indeed acquired *de novo*, and influence only the memory responses (IL-15+IL-18+LPS) and not the naive response to cytokines IL-12+IL-18 of D14LPS NK cells. We further probed the *ifng* locus for H3K4me1 after sinefungin treatment. We perfomed ChIP-qPCR on NK cells from the respective groups of mice and found that sinefungin treatment blocked H3K4 methylation at the −22kb enhancer in D14LPS - Sinefungin mice (Fig. 5e), the condition in which memory response was abrogated. Therefore, NK cell memory to endotoxemia is mediated through a histone methylation-dependent mechanism, which reveals at least one latent enhancer at the *ifng* locus.

### 6. Memory NK cells contribute to protection against bacterial infection

To investigate the potential role of endotoxemia-induced memory NK cells in the context of bacterial infection we transferred approximately 1×10^5^ highly purified D14PBS or D14LPS NK into naïve mice and infected them intraperitoneally with luminescent *E. coli*, 2 days after transfer. Bacteria were easily detected in the peritoneal cavity 15 minutes after infection (D0 in Fig. 6a) and could be followed for the next days. Noticeably, surviving mice, in which memory D14LPS NK cells were transfer, had a significantly reduced luminescent signal at day 1 post-infection compared to mice that received D14PBS NK cells (Fig. 6a,b). Memory NK cell transfer also limited mortality in recipient mice, with a 53.8% survival rate as compared to only 25% in recipients of control NK cells (Fig. 6c). Therefore, these results suggest that post-endotoxemia memory NK cells play a beneficial role in bacterial infection by contributing to a reduction in bacterial burden.

**Fig. 6.**
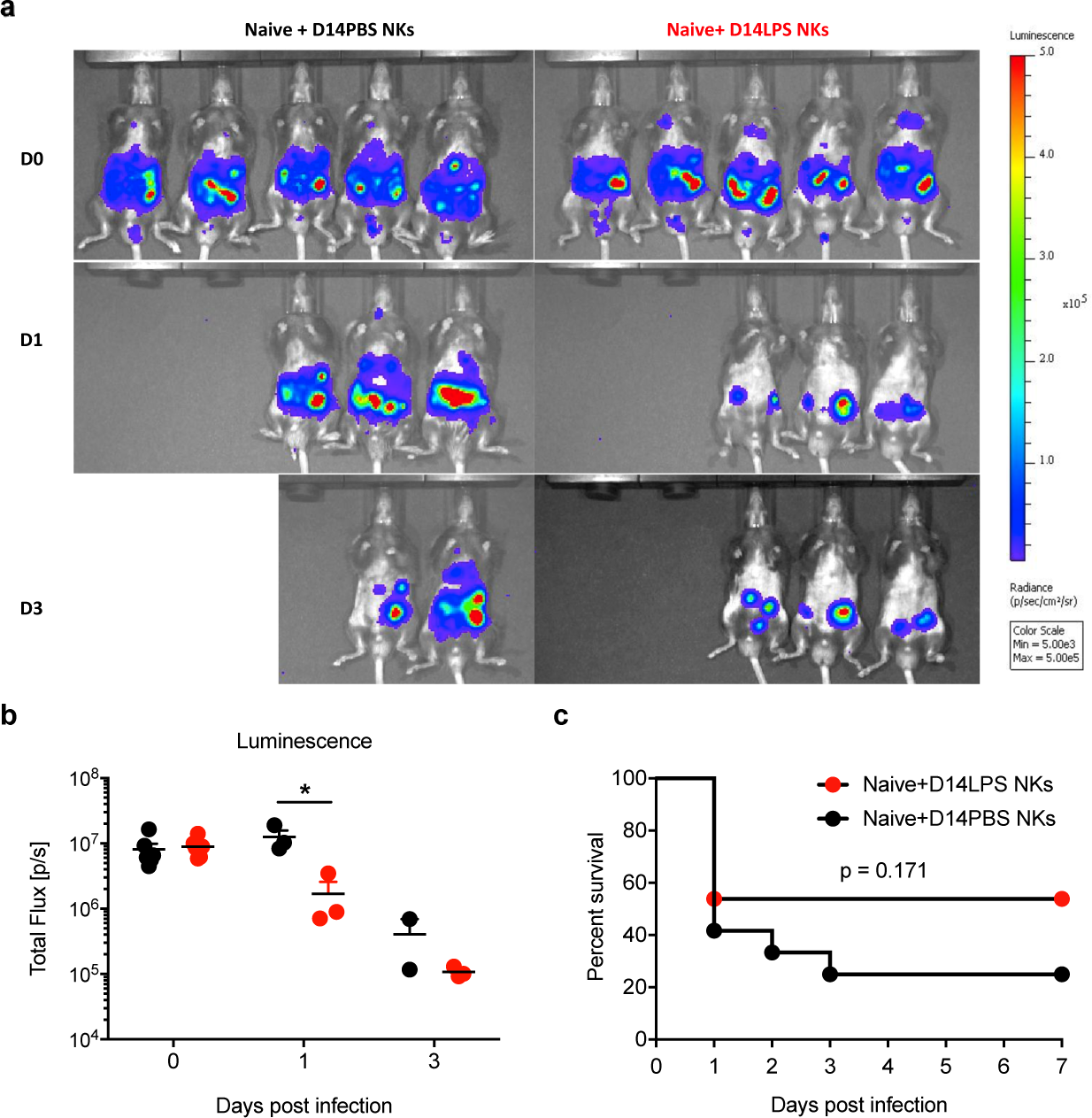
Memory NK cells protect from bacterial infection. NK cells purified from spleens of mice that were injected 14 days before with PBS (D13PBS) or LPS (10mg/kg, i.p.) (D13LPS) were transferred into naive mice (1×10^5^ cells/recipient, intravenously). Recipient mice were challenged two days later with a luminescent *E. coli* (2×10^7^CFU/mouse) via intraperitoneal injection. Luminescent signal was acquired daily for 3 days post-infection, and survival was monitored for 7 days. **a**) Representative images of bacterial luminescent signal in challenged mice at day 1 and 3. **b**) Quantification of luminescent signal. c) Survival curve for challenged mice (n=12-14/group). Data are representative of one experiment (panels a and b), or pooled results (panel c), from 2 repeats with n≥5 mice/group.

## DISCUSSION

In this study we show that LPS-induced systemic inflammation drives the formation of memory NK cells that persist for at least 9 weeks after endotoxemia, even under the control of a suppressive environment in the spleen. We demonstrate that NK memory is cell-intrinsic and is acquired by cells present at the time of systemic inflammation that undergo proliferation. We further provide molecular mechanism by which the memory is induced and/or maintained. We find that endotoxemia reveals a latent enhancer upstream of the *ifng* locus in NK cells, as seen by a dramatic increase in H3K4me1 levels, and that blocking methyltransferase activity can prevent acquisition of memory. Finally, we show that endotoxemia-induced memory NK cells can protect against bacterial infection.

Our findings that NK cells acquire memory properties, leading to enhanced activation upon re-stimulation after systemic inflammation, are surprising, since the consensus in literature is that NK cell functions are impaired during, and briefly following, acute episodes of sepsis(*13*). Indeed, when stimulated with cytokines, NK cells from acute phase sepsis patients or cecal-ligation and puncture (CLP) mice were reported to have suppressed IFNγ production(*16–18*). NK cell cytotoxicity was also shown to be impaired(*19, 33*). However, very few studies have investigated the post-acute long-term effects of systemic inflammation on NK cells and their cell intrinsic activities independent of the immune environment. A recent report suggested that NK cell responsiveness was intact 7 days after bacterial and viral pneumonia, but their activation during secondary infection was hampered by the immunosuppressive environment in the lungs(*34*). These results are similar to what we present here, where 2-9 weeks after endotoxemia, the environment restricts NK cell activation during re-stimulation, independently of their cell-intrinsic functions. Previous reports have investigated the long-term immune consequences (up to 12 weeks) of systemic inflammation in CLP mice and showed that a suppressive environment is in place, including increased Treg frequencies and their suppressive capacity(*25, 35, 36*), DC functional impairment(*24*) and expansion of immature myeloid cells(*26*). Similarly, we also observed increased frequencies of Tregs and expansion of myeloid cells, suggesting that endotoxemia induces similar long-lasting consequences to CLP. Thus, our results provide new evidence on the persistent effect of systemic inflammation on NK cell status. We show that while being within this suppressive environment, NK cell responsiveness to cytokine stimulation is not impaired. Most importantly, NK cell responsiveness to a second challenge with LPS is increased, in a cell intrinsic manner, providing the first evidence of NK cell memory following systemic inflammation.

Because of the nature of the LPS model we used in this study, a natural question is whether TLR4 is involved in endotoxemia-induced NK cell memory. Although LPS is the main ligand for TLR4, the role of this receptor for NK cells remains controversial. While several studies attributed a role for TLRs in NK cell activation(*37, 38*), TLR4 was shown to be dispensable for cell-intrinsic NK activation during endotoxemia *in vivo*, where NK cell response was mostly dependent on cytokines including IFNβ, IL-2, Il-15, IL-18 and IL-12(*39*). In line with these results, we found that activation of NK cells is not dependent on TLR2 and −4 during both primary and secondary responses *in vivo*. These results, along with our *in vitro* and *in vivo* data, where naïve and memory cells were subjected to the same inflammatory environment, argue for a minor role of cytokines in the increased responsiveness of memory NK cells. We believe that activation of NK cells during primary and secondary responses in our system would have to involve one or more NK cell receptors. Indeed, reports on NK cell memory to viruses and haptens show that specific receptors are involved in conferring a memory response(*5*). While suggested from the very first description of memory NK cells(*40*), the role of the Ly49C/I receptors in establishing NK cell memory to hapten sensitization was just recently proven(*41*). Ly49C/I – MHC-I interactions were suggested to contribute to formation of memory responses by recognizing specific amino acid sequences on peptides. Similarly, human NK cells in the context of CMV infection also recognize specific MCH-I presented peptides via the NKG2C receptor(*8*). Recognition of the m157 glycoprotein of MCVM by Ly49H(*7, 42*) and of haptens by Ly49C/I(*40, 41*) in mice, and CMV peptides by the NKG2C receptor in humans(*8*), substantiate the specificity of NK cells in both primary and recall responses. Indeed, receptor driven NK cell memory was shown to be exquisitely specific and contained within a subpopulation of cells carrying a given receptor(*8, 40, 41*). Thus, the majority of memory NK cells that have been described are characterized by a receptor, which defines memory subpopulations within the total pool. While we have excluded the role of Ly49H and NKG2A/C/E, and we have not identified a specific receptor-defined sub-population responsible for mediating post-endotoxemia memory, the possibility remains that such a population exists.

Beyond the receptors at play in NK cell memory, the molecular mechanisms involved are just beginning to be uncovered. In human and mouse memory-like NK cells, changes in DNA methylation at the *IFNG*/*ifng* locus and at various signaling molecule loci was shown to be correlated with differential transcriptional responses, suggesting that an epigenetic component could be involved(*10, 43, 44*). Recently, an extensive study of chromatin accessibility on MCMV-specific Ly49H memory NK cells using ATAC-Seq was performed(*11*). These results highlighted that memory NK possess distinct chromatin accessibility states, similarly to memory CD8 cells. However, how this regulates gene expression remains unknown. Here we show that memory NK cells acquire H3K4me1 at an upstream region of the *ifng* locus, previously identified to be crucial for IFNγ expression and suggested to be an enhancer(*31*). Such, *de novo* marking of an enhancer was observed in memory CD8 T cells following LCMV infection(*45*), but has not been reported for NK cells. In addition, H3K4me1 enhancer marking in macrophages is retained upon signal termination and is important for the augmented response observed upon re-stimulation(*46*). Interestingly, such data along with ours, strongly suggest that the enhancer repertoire of a cell is not fixed upon terminal differentiation, can be expanded by external stimuli, and can be maintained for a memory response.

NK cell memory is well described for hapten-sensitization, viral infection and *in vitro* cytokine treatment(*5*), however, evidence of NK cell memory to bacterial infection or sepsis is scarce. One report described a mouse model of BCG vaccination and *Mycobacterium tuberculosis* infection showing an expansion of IFNg+NKp46+CD27+ cells during re-infection and that transfer of these cells into naïve mice could help control bacterial burden(*47*). The authors also observed similar expansion of memory-like cells upon *in vitro* stimulation of peripheral blood mononuclear cells from humans with latent TB. NK cells from these samples could control macrophage infection with *M. tuberculosis in vitro*. We find that systemic inflammation induced by LPS also elicits NK cell memory and these memory NK cells can protect against *E. coli* infection when transferred into naïve mice. These studies and ours imply that NK memory to bacteria is relevant, and therefore memory NK cells could be considered as vaccination targets and efficacy read-outs. This prospect is especially interesting given the important role NK cells play during bacterial infection and their potency in treating such diseases as cancer (*15, 48*). In addition, our finding that NK cells responsiveness to bacterial products is heightened, while that of the environment is suppressed, suggests that NK cells could be good targets for immunotherapy in order to modulate the post-sepsis immune system.

## MATHERIALS AND METHODS

### Mice and endotoxemia model

All protocols for animal experiments were reviewed and approved by the CETEA (Comité d’Ethique pour l’Expérimentation Animale - Ethics Committee for Animal Experimentation) of the Institut Pasteur under approval number 2016-003 and were performed in accordance with national laws and institutional guidelines for animal care and use. CD45.1, CD45.2 and CD45.1/2 mice, male and females of 8 to 12 weeks were purchased from Janvier (France) or breed at the Institut Pasteur animal facility. For endotoxemia, mice were injected intraperitoneally with a single dose of conventional LPS from *E. coli* O111:B4 (Sigma Aldrich), 10 mg/kg in a 200μl volume of PBS. Control mice received just PBS injections. Animals were monitored daily for the first 5 days and then weekly for the duration of the experiment. Animals were sacrificed in the event they lost more than 20% of weight as a result of endotoxemia (mortality was usually below 10% at this dose of LPS). A clinical score was used to assess progression of mice throughout endotoxemia, representing the sum of the following scores: 0 – no clinical signs; 1 – hypoactivity; 1 – ruffled fur; 2 –hunched posture; 2 – diarrhoea; 3 – prostration. For histone methyltransferase inhibition experiments, mice were treat at day 2 after PBS/LPS injection with saline or Sinefungin (Abcam) (10 mg/kg, i.p. in a 200μl volume of saline.

### Cytometry staining protocols and analysis

Single cell suspensions from spleens were counted and prepared for surface staining in 96 well plates. Unspecific binding was first blocked by incubation with anti-mouse CD16/CD32 (BD Biosciences) for 10 minutes before addition of surface labelling antibodies for another 45 minutes, in 0,5% FCS at 4°C. Cells were washed twice in PBS in preparation for viability staining using fixable viability dye (eFluor780, ebioscience) for 5 minutes at 4°C. Cell were washed twice in 0,5% FCS and fixed using commercial fixation buffer (Biolegend). For intracellular cytokine and Foxp3 staining, cells were permeabilized and washed with buffers from commercial kits (Inside Stain Kit, Miltenyi Biotec and Foxp3 / Transcription Factor Fixation/Permeabilization Concentrate and Diluent, eBioscience, respectively). Following permeabilization and washing, cells were stained with respective antibodies in 0,5% FCS for 30 minutes at 4°C and after a final washing suspended in buffer for FACS analysis. Sample acquisition was performed on a MACSQuant flow-cytometer (Miltenyi Biotec) and analysis was done using FlowJo Software (TreeStar).

### >Antibodies

The following antibodies (clones) were used: NK1.1 (PK136), CD3 (145-2C11), CD49b (DX5), CD69 (H1.2F3), Ly6C (HK1.4), Ly6G (1A8-Ly6g), CD11b (M1/70), CD11c (HL3), Ly49H (REA241), Ly49F (HBF-719), Ly49D (4E5), Ly49C/I (REA253), NKG2A/C/E (20d5), NKG2D (CX5), CD45.1 (A20), CD45.2 (104), foxp3 (FJK-16s) and IFNγ (XMG1.2) were purchased from Biolegend, eBioscience, BD Biosciences or Miltenyi Biotec.

### NK cell enrichment and purification

Splenocytes obtained by mechanical dissociation of spleens were passed successively through 100μm, 70μm and 30μm strainers (Milteny Biotec) and counted before use for downstream applications. NK cells were enriched from splenocyte suspensions using negative enrichment kits (eBioscience) according to manufacturer’s protocol but combined with separation over magnetic columns (LS columns, Miltenyi Biotec) and were routinely brought to 80% purity. For some *in vitro* stimulations and transfers of cells before bacterial challenge, enriched NK cells were re-purified using mouse NK cell purification kits (Miltenyi Biotec) according to manufacturer’s instructions to reach purities of approximately 98%.

### Cell transfer experiments

Enriched NK cells from CD45.1, CD45.2 or CD45.1/2 mice were transferred i.v. into recipient congenic mice (0,75-1×10^6^ cells/recipient) in 100μl of PBS. For forward transfers, NK cells were transferred into congenic naïve mice 1 day before LPS challenge. At 6 hours after PBS or LPS injection, single cell suspensions were obtained from spleen of recipient mice and flow cytometric analysis was performed. For pre-/post-transfer experiments, CD45.1 NK cells were pre-transferred into CD45.1/2 mice 1 day before endotoxemia induction and naïve CD45.2 NK cells were post-transferred 7 days after. At day 14 after endotoxemia, mice were re-challenged with PBS/LPS and NK cell responses were assessed by flow cytometry as described above. For proliferation assessment experiments, enriched NK cells were stained with CFSE 2,5μM in PBS for 10 minutes, before blocking in FCS and extensively washed with PBS before transfer into congenic mice, 1 day before endotoxemia induction. For bacterial challenge experiments 10^5^ highly-purified NK cells from D13PBS or LPS mice were transferred to naïve 2 days before challenge with 2-4×10^7^ *E. coli* intraperitoneally.

### NK cell culture and co-cultures

Purified NK cells were cultured at 1×10^6^ cells/ml in 200μl RPMI 1640 (Gibco) supplemented with 10% FCS and Penicillin-Streptomycin (Gibco) respectively in 96 well round-bottom plates (Nunc). Cells were either left unstimulated or activated with conventional LPS from E. coli O111:B4 (Sigma Aldrich) at 100ng/ml or various cytokine cocktails composed of IL-15, IL-12 (Miltenyi Biotec) and IL-18 (MBL International) at a dose of 10ng/ml each. NK cell were additionally stimulated with LPS + cytokine cocktails. After 20 hours of culture, cells were collected for intracellular cytokine staining by flow cytometry and culture supernatants were collected and stored at −20°C for cytokine assays performed by ELISA according to manufacturer’s instructions (DuoSet, R&D Systems).

### ChIP-qPCR assays

ChIP was performed as described before by Batsche *et al*(*49*) with some modifications. NK cells were fixed in 1% formaldehyde (8 min, room temperature), and the reaction was stopped by the addition of glycine at the final concentration of 0,125 M. After two washes in PBS, cells were resuspended in 0.25% Triton X-100, 10 mM Tris-HCl (pH 8), 10 mM EDTA, 0.5 mM EGTA and proteases inhibitors; the soluble fraction was eliminated by centrifugation; and chromatin was extracted with 250 mM NaCl, 50 mM Tris-HCl (pH 8), 1 mM EDTA, 0.5 mM EGTA and proteases inhibitors cocktail for 30 min on ice. Chromatin was resuspended in 1% SDS, 10 mM Tris-HCl (pH 8), 1 mM EDTA, 0.5 mM EGTA and proteases inhibitors cocktail; and sonicated during 6 cycles using Diagenode Bioruptor Pico (15 sec on, 30 sec off). DNA fragment size (<1 kb) was verified by agarose gel electrophoresis. ChIP was performed using H3, H3K4me3, H3K4me1 antibodies and nonimmune IgG (negative control antibody), chromatin extracted from 3 x 10^6^ NK cells per condition was used. Chromatin was diluted 10 times in 0.6% Triton X-100, 0.06% sodium deoxycholate (NaDOC), 150 mM NaCl, 12 mM Tris-HCl, 1 mM EDTA, 0.5 mM EGTA and proteases inhibitors cocktail. For 6 hours, the different antibodies were previously incubated at 4°C with protein G-coated magnetic beads (DiaMag, Diagenode), protease inhibitor cocktail and 0.1% BSA. Chromatin was incubated overnight at 4°C with each antibody/protein G-coated magnetic beads. Immunocomplexes were washed with 1 x buffer 1 (1% Triton X-100, 0.1% NaDOC, 150 mM NaCl, 10 mM Tris-HCl (pH 8)), 1 x buffer 2 (0.5% NP-40, 0.5% Triton X-100, 0.5 NaDOC, 150 mM NaCl, 10 mM Tris-HCl (pH 8)), 1 x buffer 3 (0.7% Triton X-100, 0.1% NaDOC, 250 mM NaCl, 10 mM Tris-HCl (pH 8)) 1 x buffer 4 (0.5% NP-40, 0.5% NaDOC, 250 mM LiCl, 20 mM Tris-HCl (pH 8), 1 mM EDTA), 1 x buffer 5 (0,1% NP-40, 150 mM NaCl, 20 mM Tris-HCl (pH 8), 1 mM EDTA) and 1 x buffer 6 (20mM Tris-HCl (pH 8), 1 mM EDTA). Beads were eluted in water containing 10% Chelex and reverse cross-linked by boiling for 10 min, incubating with RNase for 10 min at room temperature, then with proteinase K for 20 min at 55°C and reboiling for 10 min. DNA fragment were purified by Phenol Chloroform extraction.

Amplifications (40 cycles) were performed using quantitative real-time PCR using iTaq^TM^ Universal Syber Green Supermix (BIORAD) on a CFX384 Touch Real-Time PCR system (BIORAD). qPCR efficiency (E) was determined for the ChIP primers with a dilution series of genomic mouse DNA. The threshold cycles (Ct values) were recordered from the exponential phase of the qPCR for IP and input DNA for each primer pair. The relative amont of immunoprecipitated DNA was comparated to input DNA for the control regions (% of recovery) using the following formula: % recovery = E^((Ct(1% input) - Log_2_(input dilution)) – Ct (IP)) x 100. Antibodies ChIP-grade and ChIP primers sequences are provided in Supplemental Table 1

### Statistical analysis

Statistical significance was tested using Prism 5.0 Software (GraphPad). Mann Whitney test was used for single comparisons and Wilcoxon paired test for paired comparison of samples in the same mouse in transfer experiments or in the same well for co-culture experiments. Unless otherwise specified in figure legends, error bars in all figures represent SEM, with the midlines representing the mean value.

## ACKNOWLEDGEMENTS

Work in the M.A.H. laboratory received financial support from Institut Pasteur and the National Research Agency (ANR-EPIBACTIN). O.R. was supported by a stipend from the Pasteur - Paris University International Ph.D. Program for part of this work. T.C. is supported by a fellowship from the Foundation for Medical Research (Marianne Josso Prize).

## CONTRIBUTIONS

O.R. coordinated the study; O.R., J-M.C. and M.A.H. conceived of and designed the study; O.R., T.C., C.C. and C.F. performed experiments and analyzed data; O.R. and M.A.H. conceived and wrote the manuscript; J-M.C. edited the manuscript; J-M.C. and M.A.H. supervised the work

---

**Fig. S1.**
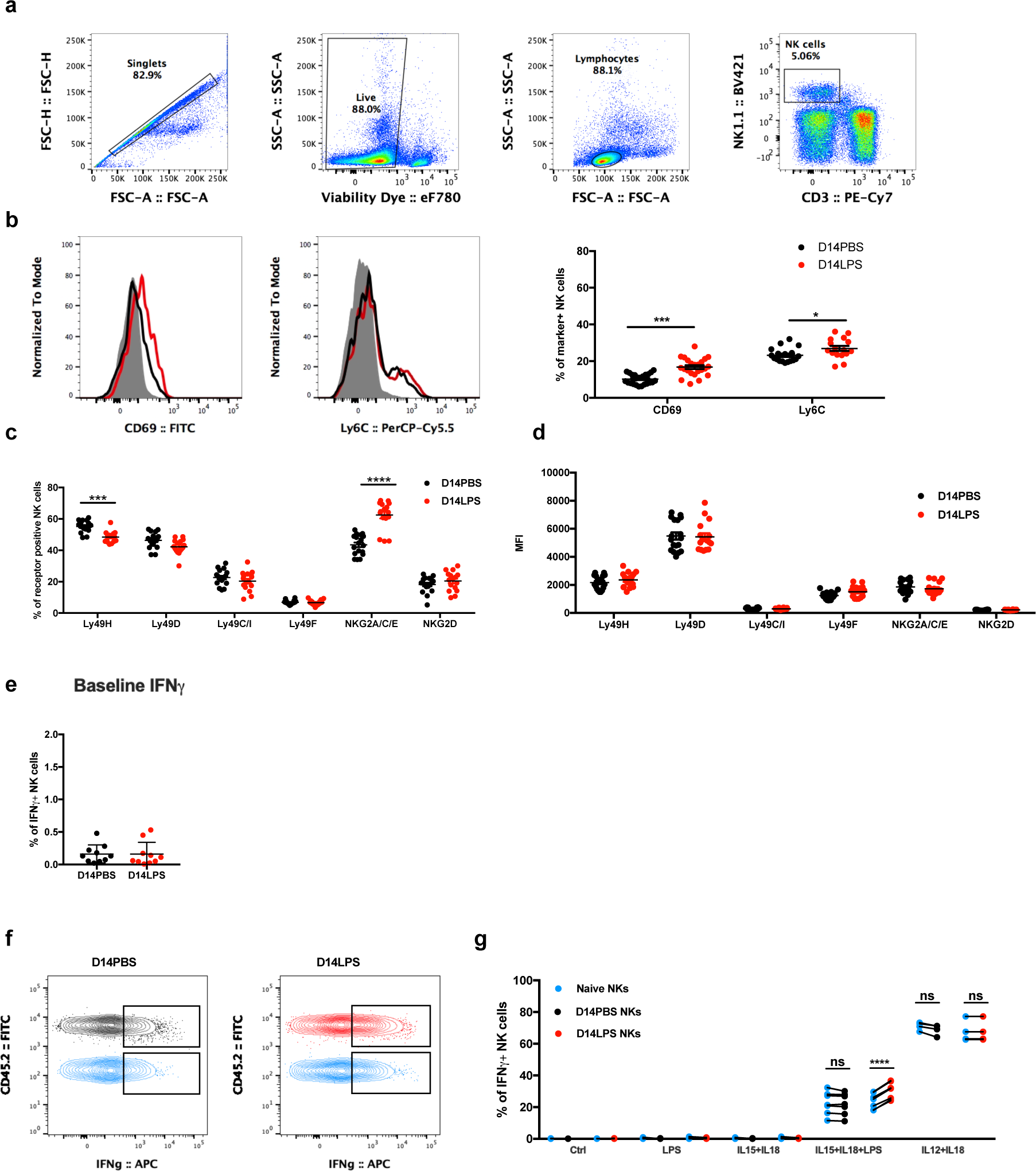
NK cell gating strategy and NK receptor profiles after endotoxemia. Splenocytes were harvested from mice 2 weeks after they were intraperitoneally injected with PBS (D14PBS) or LPS (10mg/kg, i.p.) (D14LPS) and stained for NK cell identification (CD3-NK1.1+) and NK cell receptors. **a)** Gating strategy used for NK cell identification. **b)** representative overlay histograms and quantification of CD69 and Ly6C expression **c)** Data summary of NK cell expression of several NK cell receptors in percentages and **d)** intensity of expression. **e)** Quantification of *ex-vivo* IFNγ staining. Higly purified CD45.2 D14PBS or D14LPS NKs were mixed with enriched naïve CD45.1 NKs and stimulated overnight in the indicated conditions. Cells were stained for flow cytometry analysis of intracellular IFNγ levels. **f)** Representative contour plots of IFNg staining among the co-cultured NKs. **g)** Data summary grouped before-after plots depicting percentages of IFNγ+ cells among CD45.1 naïve (blue dots), CD45.2 D14PBS (black dots) and D14LPS (red dots) NKs. Dots represent individual mice except for panels e,f where they represent technical repeats from group-pooled samples. Horizontal lines represent mean values +/-SEM. Data represent pooled values from 2-5 repeats with n≥ 5 mice/group for b and c, and one representative experiment out of 3 repeats with n≥5 mice/group for panels e.f. ns, not significant, * p<0.05, ***p<0.001. Mann–Whitney test comparing D14Sln and D14LPS values under respective conditions.

**Fig. S2.**
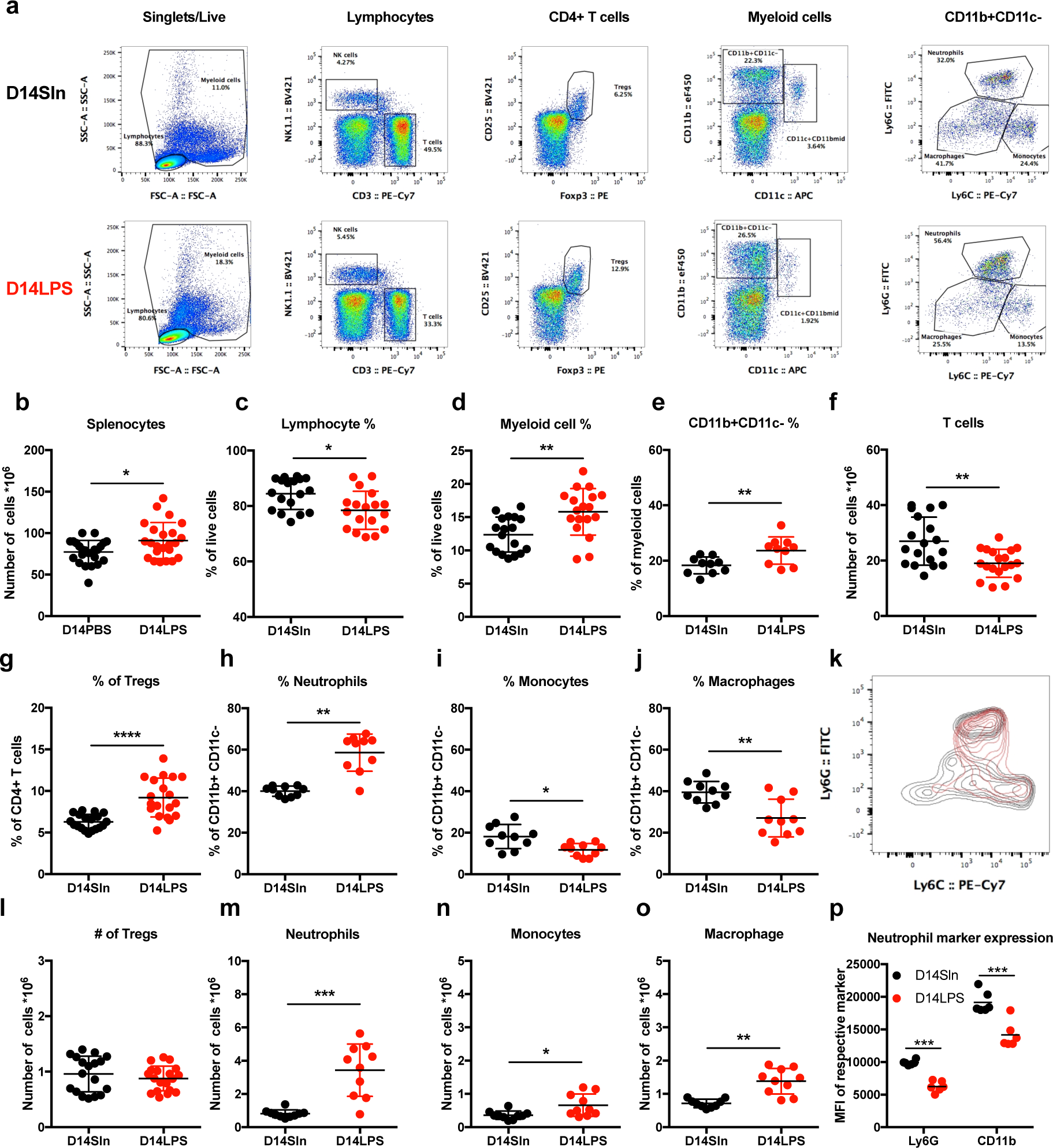
Spleen environment is altered after endotoxemia. Splenocytes were harvested from mice 2 weeks after they were intraperitoneally injected with PBS (D14PBS) or LPS (10mg/kg, i.p.) (D14LPS). Cells were stained for identification of T cells (NK1.1-CD3^+^), NK cells (NK1.1^+^CD3), Tregs (CD3^+^CD4^+^CD25^+^Foxp3^+^), neutrophils (CD11b^+^CD11c-Ly6G^high^Ly6C^+^), monocytes (CD11b^+^CD11c-Ly6G-Ly6C^high^) and macrophages (CD11b^+^CD11c-Ly6G^-/+^Ly6C^-/+^). (a) Representative dot plots depicting gating strategy and sample results from D14PBS and D14LPS splenocyte staining. (b) Total splenocyte numbers. (c) Lymphocyte and (d) myeloid cell percentages among live events. (e) Percentage of CD11b+CD11c-cells among myeloid cells. (f) Total T cell numbers. (g) Percentage and (l) total number of Tregs. (h) Percentage and (m) total number of neutrophils. (i) Percentage and (n) total number of monocyte. (j) Percentage and (o) total number of macrophages. (k) Representative overlay contour plot of Ly6G/Ly6C profiles of myeloid cells from D14PBS (black) and D14LPS (red). (p) Quantification of staining intensity of CD11b and Ly6G on neutrophils. Horizontal lines represent mean values +/-SEM. Dots represent individual mice. Data represent pooled values from 2-5 repeats with n≥ 5 mice/group ns, not significant, * p<0.05, ***p<0.001. Mann–Whitney test comparing D14PBS and D14LPS values under respective conditions.

**Fig. S3.**
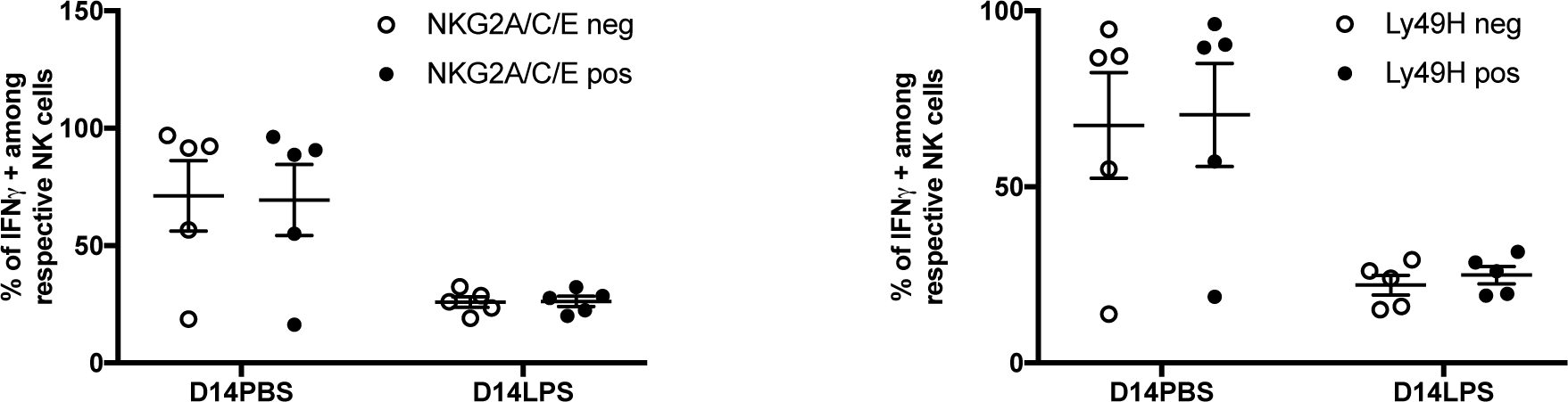
NK cell response during LPS challenge in different sub-populations. Mice that were intraperitoneally injected 14 days in advance with PBS (D14PBS) or LPS (10mg/kg, i.p.) (D14LPS) were rechallenged with LPS (10mg/kg, i.p.). Six hours after re-challenge, splenocytes were processed for flow cytometry to identify NK cells positive and negative for NKG2A/C/E or Ly49H and the levels of intracellular IFNγ in the respective sub-populations. **a**) Data summary of IFNγ+ percentages amonge the respective NKG2A/C/E positive or negative populations in D14PBS and LPS mice. **b**) Data summary of IFNγ+ percentages amonge the respective Ly49H positive or negative populations in D14PBS and LPS mice. Bars represent mean values +/-SEM. Dots represent individual mice. Data are representive of one out of 2 repeats n=5 mice/group ns, not significant. Mann–Whitney test comparing percentages of IFNγ+ among NKG2A/C/E or Ly49H positive to negative populations under respective conditions.

**Fig. S4:**
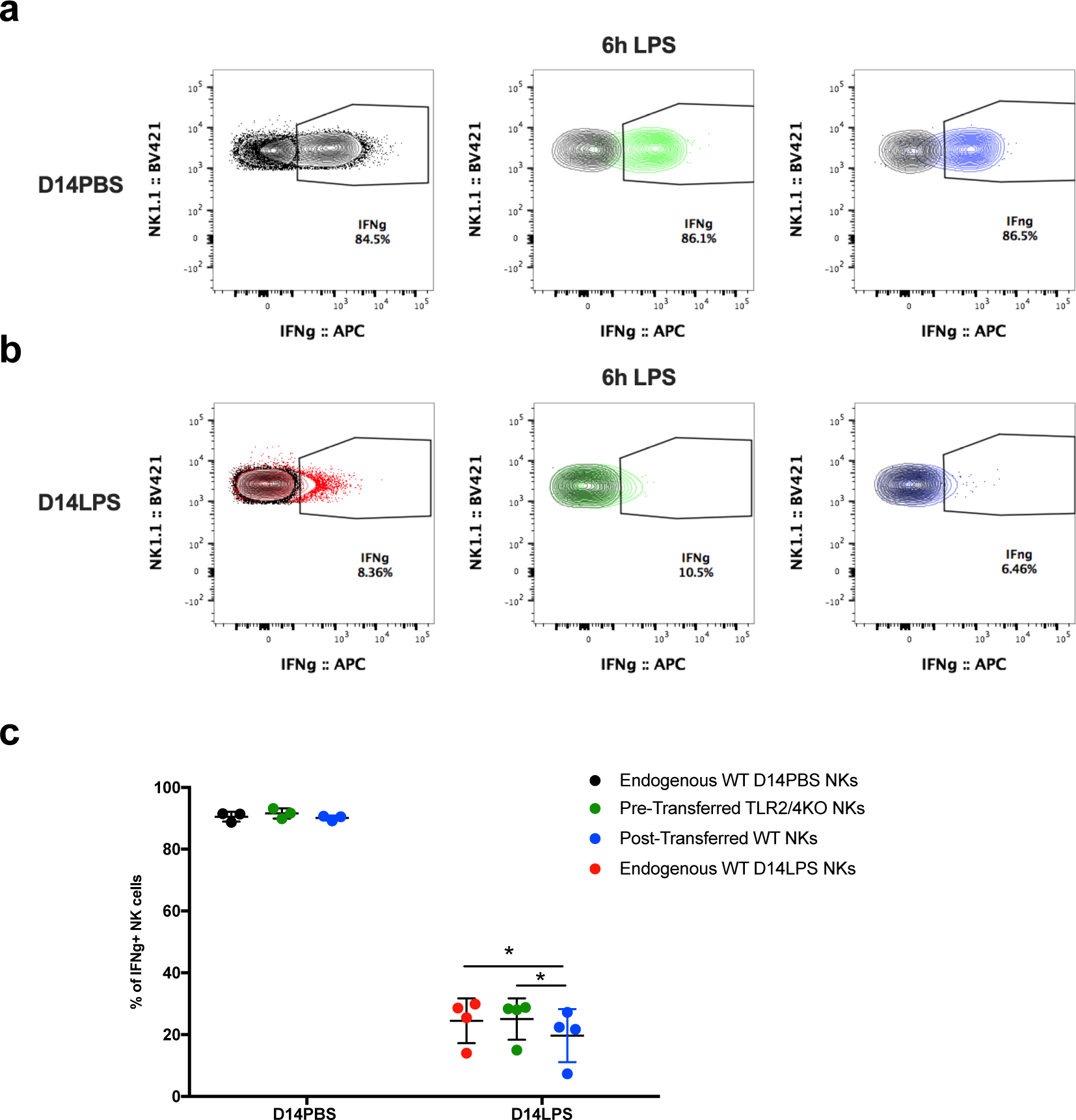
NK cell responses in pre-post transfers. NK cells enriched from spleens of CD45.1 mice were Pre-Transferred into CD45.1/2 mice (1×10^6^ cells/recipient, intravenously), one day before recipients were injected with PBS or LPS (10mg/kg, i.p.). Naïve CD45.2 enriched NK cells were Post-Transferred into the CD45.1/2 mice, 7 days after PBS/LPS injection. On day 14 with CD45.1/2 D14PBS and D14LPS mice were rechallenged with LPS and sacrificed 6 hours later to assess intracellular IFNγ levels in endogenous, pre- and post-transferred NK cells. **a)** Representative results depicting percentages of IFNγ+ cells among endogenous CD45.1/2 D14PBS NKs (black), Pre-Transferred CD45.1 NK cells (green), Post-Transferred CD45.2 (blue) **b)** and endogenous CD45.1/2 D14LPS NK cells (red) at 6 hours after LPS rechallenge. NK cells enriched from spleens of CD45.2 TLR2/4KO mice were Pre-Transferred into WT CD45.1/2 mice (1×10^6^ cells/recipient, intravenously), one day before recipients were injected with PBS or LPS (10mg/kg, i.p.). Naïve CD45.1 enriched NK cells were Post-Transferred into the CD45.1/2 mice, 7 days after PBS/LPS injection. On day 14 with WT CD45.1/2 D14PBS and D14LPS mice were rechallenged with LPS and sacrificed 6 hours later to assess intracellular IFNγ levels in endogenous, pre-and post-transferred NK cells. **c)** Data summary depicting percentages of IFNγ+ cells among endogenous WT CD45.1/2 D14PBS NKs (black dots), Pre-Transferred TLR2/4KO CD45.2 NK cells (green dots), Post-Transferred WT CD45.1 (blue dots) and endogenous WT CD45.1/2 D14LPS NK cells (red dots) at 6 hours after LPS rechallenge. Dots represent individual mice. Data are representative of one experiment out of 2 repeats with n=3-4mice/group. ns, not significant, * p<0.05. Wilcoxon paired test comparing Pre, Post and Endogenous NK cells under respective conditions.

**Table S1.**
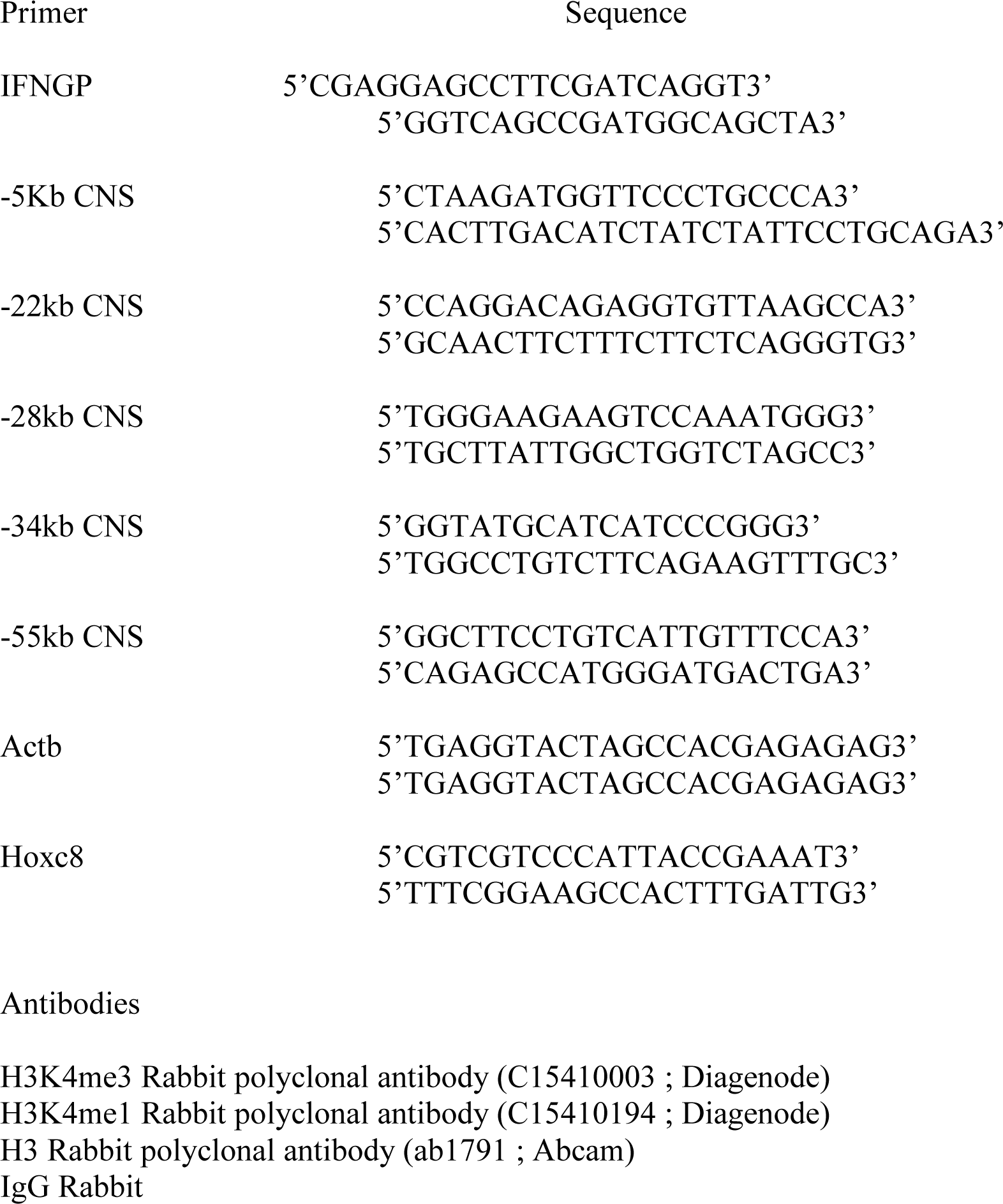
ChIP primer sequences and antibodies

